# Neutrophil reverse migration from liver fuels neutrophilic inflammation to tissue injury in Nonalcoholic Steatohepatitis

**DOI:** 10.1101/2021.10.03.462893

**Authors:** Maria Feliz-Norberto, Cassia Michael, Sofia de Oliveira

## Abstract

Inflammation is a hallmark in the progression of nonalcoholic-fatty liver disease (NAFLD) to non-alcoholic steatohepatitis (NASH). Patients with NAFLD are characterized by a chronic low-grade systemic metabolic inflammation (i.e., metainflammation), which contributes to exacerbated however dysfunctional immune response. Neutrophils play an important pathological role in NAFLD progression to NASH; however, how NASH and associated chronic systemic inflammation impact overall the neutrophil response to injury is completely unexplored. Here, we investigated how neutrophil response to tissue injury is altered by the presence of NASH. We used a diet-induced NASH zebrafish model combined with tailfin transection in transgenic zebrafish larvae to study neutrophilic inflammation. Live non-invasive confocal microscopy was used to investigate neutrophil recruitment to tailfin injury through time. Photoconvertion of neutrophils at the liver area followed by time-lapse microscopy was performed to evaluate migration of neutrophils from liver to tailfin injury. Metformin and Pentoxifylline were used to pharmacologically reduce NASH and liver inflammation. We found that larvae with NASH display systemic inflammation and increased myelopoiesis. NASH larvae display a dysfunctional and exacerbated neutrophil response to tailfin injury, characterized by increased neutrophil recruitment, and delayed resolution of inflammation. Interestingly, we showed that neutrophils undergo reverse migration from the NASH liver to the wounded tailfin area. Finally, pharmacological treatment of NASH with Pentoxifylline and Metformin significantly reduced systemic chronic inflammation and the exacerbated recruitment of neutrophils to tissue injury. Taken together, our findings suggest that NASH exacerbates neutrophilic inflammation probably via neutrophil priming at the liver, which can further undergo reverse migration and respond to secondary inflammatory triggers such as tissue injury. Reverse migration of primed neutrophils from the liver might be an important mechanism that fuels the exacerbated neutrophil response observed in NASH conditions and associated metainflammation contributing to poor prognosis and increasing death in patients with metabolic syndrome.

## Introduction

Nonalcoholic fatty liver disease (NAFLD) is the hepatic manifestation of metabolic syndrome (i.e., high blood pressure, high blood sugar, excess body fat around the waist, and abnormal cholesterol levels) and is a major health issue and economic burden in western societies affecting around 25-30% of overall population ^1, 2^. The rising incidence of NAFLD correlates strongly with the prevalence of metabolic syndrome, type 2 diabetes and obesity^1^. Consumption of calorie rich diets drastically increase fatty acid availability that causes local and systemic metabolic alterations and steatosis, hepatocyte injury, inflammation, and fibrosis, all of which are key features of nonalcoholic steatohepatitis (NASH) a more advanced stage of NAFLD ^3^. Systemic metabolic dysfunction triggers a chronic systemic low-grade of inflammation (i.e., metainflammation), generating immune imbalances, from cellular to cytokine levels that predisposes patients with NASH and associated metabolic diseases to chronic inflammation and infections^4, 5^. The high incidence of such complications drastically impacts this high-risk group, both socially and economically, and often leads to disability or death - as shown recently with COVID-19 pandemic^6, 7^. The lack of efficient therapeutic approaches to decrease such impact in this high-risk population is a clear indicator that we do not fully understand how metabolic syndrome, nutrient excess or overnutrition, and associated metainflammation are altering and regulating the overall inflammatory response towards “secondary” inflammatory triggers such as tissue injury.

Neutrophils are first line responders to injury that rely in distinct tiers of arsenals to counter threats including phagocytosis, protease secretion, and neutrophil extracellular traps (NETs)^8^. Such mechanisms are not only protective but can also be destructive to tissues, therefore neutrophil production, trafficking, and clearance need to be tightly regulated ^9^. Neutrophils have a double-edge sword function being crucial for effective tissue repair but can also contribute for further damaged in case a dysfunctional response is triggered^8^. Neutrophils have a crucial role on NAFLD pathophysiology; with circulating neutrophils from patients with NASH exhibiting an activated and immunosuppressive phenotype^10^. Multiple reports have also found that circulating neutrophils in NASH have enhanced reactive oxygen species (ROS) production upon inflammatory stimulus, and undergo spontaneous NETs formation (i.e., NETosis)^10-12^, based on this evidence we decided to explore how neutrophilic inflammation to tissue injury was impacted in a NASH background.

The small vertebrate animal model, the zebrafish, with its unparallel transparency and genetic similarity with humans provides a unique opportunity to explore neutrophilic inflammation in a whole-animal context using non-invasive live imaging^13^. Here, we used a diet-induced NASH zebrafish model ^14, 15^ by exposing transgenic zebrafish larvae with fluorescently-tagged neutrophils to a high cholesterol diet for one week and performed tailfin transection ^16^. Next, we investigated neutrophil recruitment to tailfin injury by live non-invasive confocal microscopy. We demonstrated that zebrafish larvae with NASH have systemic chronic inflammation, and increased myelopoiesis, which resulted in increased number of neutrophils and macrophages. Importantly, we found that NASH larvae have an exacerbated neutrophil response to tailfin injury, characterized by increased neutrophil recruitment and delayed resolution of inflammation. We also found that neutrophils undergo reverse migration from the NASH liver to the tailfin injury. Finally, we demonstrated that pharmacological treatment of NASH with Pentoxifylline and Metformin significantly reduced systemic chronic inflammation and the exacerbated recruitment of neutrophils to tissue injury. Our findings suggest that NASH exacerbates neutrophilic inflammation probably via neutrophil priming at the liver, which can further undergo reverse migration and respond to secondary inflammatory triggers such as tissue injury.

## Material and Methods

### Zebrafish general procedures

All protocols using zebrafish in this study were approved by the University of Wisconsin-Madison and Albert Einstein College of Medicine Institutional Animal Care and Use Committees (IACUC). Adult zebrafish and embryos up to 5 days post-fertilization (dpf) were maintained as described previously ^17^. At 5 dpf, larvae were transferred to feeding containers and kept in E3 embryo medium (E3) [5mM NaCl (Fisher Scientific), 0.17 mM KCl (Dot Scientific), 0.33 mM CaCl_2_ (Acros Organics), 0.33 mM MgSO_4_•7x H_2_O (Sigma-Aldrich)] without methylene blue, until the end of the experiment. For all experiments, larvae were anesthetized in E3 media without methylene blue supplemented with 0.16 mg/ml Tricaine (MS222/ethyl 3-aminobenzoate; Sigma-Aldrich-Aldrich).

### NASH zebrafish model

Larvae diets were prepared as previously described ^14, 15, 18^ using Golden Pearl Diet 5-50 nm - Active Spheres (Brine Shrimp Direct). At 5 days post fertilization (dpf), zebrafish larvae were separated into treatment groups in E3 without methylene blue as described before^14^. Briefly, 5 dpf larvae were separated into different feeding tanks corresponding to normal diet (ND) and 10% high cholesterol diet (HCD) and fed 0.1 mg of food per larvae per day. In general, 60-80 larvae were placed in a breeding container with 400 mL of E3 without methylene blue and fed 6-8 mg of ND or HFD daily. Feeding boxes were cleaned and E3 was replaced daily. Short-term feeding was performed from 5 to 12 dpf. Before experimental procedure, larvae were fasted for 18 hours to decrease intestine autofluorescence. At 13 dpf, larvae were anesthetized and screened for neutrophil and macrophage markers as needed on Zeiss Axio Zoom stereo microscope (EMS3/SyCoP3; Zeiss; Zeiss; PlanNeoFluar Z 1X:0.25 FWD 56mm lens).

### Transgenic zebrafish lines

Double transgenic line expressing human fluorescently-tagged histone-2b (H2B) in macrophages (mpeg1 promoter) and neutrophils (lyzC promoter)^19, 20^ Tg(mpeg1:H2B-EGFP; lyzc:H2B-mCherry) ^uwm43Tg/uwm40Tg^ ^19, 21^, were used for all experiments with the exception of EDU assay, Cell ROX assay, and photoconversion assay where wild-type, Tg(mpx:mCherry) ^uwm7Tg^ ^22^ and Tg(mpx:dendra)^uwm4Tg^ ^23^ were used respectively.

### Fixation of larvae

All larvae were fixed in 2 mL round bottom tubes with 2 mL of fixation solution [1.5% Formaldehyde (Polysciences, Inc.), 0.1 M PIPES (Sigma-Aldrich), 1 mM MgSO_4_ (Sigma-Aldrich), 2 mM EGTA (Sigma-Aldrich)] overnight at 4 °C. The next day larvae were rinsed and washed once for 5 min in PBS pH 7.4 (Sigma-Aldrich) and stored at 4° C in PBS until imaging ^14^.

### Confocal Microscopy Imaging-zWEDGI

All imaging was performed using a zWEDGI device as previously described ^24^. Briefly, an anesthetized larva was loaded into a zWEDGI chamber for time-lapse imaging. The loading chamber was filled with 1% low melting point agarose (Sigma-Aldrich) in E3 to retain the larvae in the proper position. Additional E3 supplemented with 0.16 mg/ml Tricaine was added as needed to avoid dryness and provide required moisture to zebrafish larvae during imaging acquisition. All images were acquired on a spinning disk confocal microscope (CSU-X; Yokogawa) with a confocal scanhead on a Zeiss Observer Z.1 inverted microscope equipped with a Photometrics Evolve EMCCD camera, and an EC Plan Neofluar NA 0.3/10 x air objective, z-stacks, 5 μm optical sections and 512 × 512 resolution. For whole-larvae imaging, 7 × 1 tile images were taken and automatically stitched. For time-lapse movies of tailfin injury and neutrophil and macrophage chemotaxis, images were taken every 2 minutes up to 16 hours post-wounding. For photoconversion assay, images were taken every 15 minutes up to 6 hours post-wounding. For Cell ROX imaging, NA 0.5/20 x air objective was used to acquire tailfin images.

### EDU incorporation, labeling, and quantification

Proliferation in whole larvae was measured using EDU staining, larvae were incubated in 10 μM 5-ethynyl-2’-deoxyuridine (EdU) dissolved in embryo medium for 6 hours. Larvae were euthanized and fixed in 4% Paraformaldehyde (Sigma-Aldrich) overnight at 4°C and stored in Methanol at -20°C until staining. Click-iT EdU Imaging Kit (Life Technologies) were used for staining following manufacturer’s instructions. Whole-larvae images were acquired as described previously. For quantification, whole-larvae images acquired and analyzed using IMARIS Bitplane software (Version 9.5/9.6) rendering mode. The number of EDU positive cells were automatically counted in whole larvae using the IMARIS spots function. Spots were defined as particles with 5 μm and 10 μm of X/Y and Z diameter, respectively. To quantify the number of EDU positive cells at the hematopoietic niches such as, caudal hematopoietic tissues (CHT), kidney, and thymus a surface for each niche was created. Then a mask for the EDU signal was generated to isolate the signal for EDU positive cells at the hematopoietic niches. EDU positive cells were automatically counted using IMARIS spots function and same parameters as for whole larvae.

### Tailfin transection

At 13 dpf, transgenic larvae fed with normal or high cholesterol diet were transferred from feeding boxes into petri dishes with fresh E3 without methylene blue. Next, larvae were anesthetized in E3 with 0.16 mg/ml of tricaine. Complete transection of the tailfin tip was then performed using a scalpel with a n°10 sterile surgical blade under a stereomicroscope equipped with a transillumination base (Nikon SMZ 745; Nikon) without harming the notochord. The success of transection was immediately confirmed. Larvae were then transferred to a new plate with E3 without methylene blue and let to recover at 28°C until collection timepoint or were mounted immediately in a zWEDGI for time-lapse confocal microscopy imaging. In photoconversion assay, tailfin transection was performed with larvae mounted in the zWEDGI.

### Automatic quantification of innate immune cells and cell-tracking on IMARIS

To quantify number of neutrophils and macrophages 13 dpf larvae were fixed and whole-larvae or tailfin images were acquired as described previously. For quantification of the number of neutrophils and macrophages in whole-larvae or at different regions, acquired whole-larvae images were reconstructed on IMARIS Bitplane software (Version 9.5/9.6) rendering mode and total number of neutrophils and macrophages were automatically counted in the whole larvae using IMARIS spots function. Spots were defined as particles with 5 μm and 10 μm of X/Y and Z diameter, respectively. To quantify the number of neutrophils and macrophages at different regions of the zebrafish larvae, a surface for each region/area was created, then a mask for the neutrophil and macrophage signals were generated setting to “zero” signal outside the surface. Cells were automatically counted using the IMARIS spots function and defined as particles with 5 μm and 10 μm of X/Y and Z diameter, respectively. To quantify neutrophil recruitment to a tailfin wound acquired time-lapse movies or images of transected tailfins were reconstructed on IMARIS software. Neutrophil recruitment was assessed at wound sites (the region posterior to the circulatory loop, Figure 3A) at various time points (1-, 2-, 4-, 6- and 8-hours post wounding) for the neutrophil recruitment time-course and 4 and 24hpw for the resolution experiments. Finally, acquired time-lapse movies (8h acquisition) were used to perform automatic neutrophil tracking to the tailfin transection on IMARIS. Cell tracking was performed in the field of view (FOV) and in the wounded area (the region posterior to the circulatory loop, Figure 3A). Mean neutrophil speed was obtained by IMARIS cell tracking analysis. IMARIS volume rendering mode was used to obtained representative 3D reconstructions that were used for figures and supplemental movies.

### Systemic Chronic Inflammation (SCI) and neutrophil CHT depletion scorings

To address the systemic effect of cholesterol-enriched diet on neutrophils and macrophages 13 dpf transgenic larvae were fixed and whole-larvae images were acquired as described previously. Acquired whole-larvae images were analyzed using IMARIS rendering. Larvae from different diets or treatments were scored in two different ways, one for systemic chronic inflammation (SCI) and a second score to evaluate neutrophil depletion from caudal hematopoietic tissue (CHT). For such, infiltration of neutrophils and macrophages to tissues and organs as well as reduction of the number of neutrophils from CHT were performed in each larva. Scoring for No SCI, Mild SCI or Severe SCI and for Normal, Mild depletion or Severe depletion/ empty CHT, as shown in representative images in Suppl. Figure 3.

### Resolution Assay

To address if resolution of inflammation was occurring, tailfin transection was performed on anesthetized 13 dpf larvae fed with ND or HCD, then larvae were fixed at 4 and 24 hours-post wounding (hpw). Tailfin images were acquired and automatic quantification of number of neutrophils recruited to wound area at 4 hpw (peak of recruitment) and 24hpw (resolution) was performed as described using IMARIS Bitplane Software.

### Cell ROX assay and quantification

To measure ROS production by neutrophils, tailfin transection was performed on anesthetized 13 dpf Tg(mpx:mCherry) larvae fed with ND or HCD as previously described. Larvae were let to recover in E3 for 3 hours and 30 minutes and then incubated for 45 minutes in 5 µM CellROX® Deep Green Reagent (Invitrogen) solution diluted in E3 without methylene blue. Larvae were fixed and tailfin images were acquired as described. To address ROS production in neutrophils, acquired images were analyzed using IMARIS (Bitplane) rendering, IMARIS Coloc mode was used to quantify number of colocalized voxels and percentage of dataset colocalized in tailfin wound using neutrophil and Cell ROX signals.

### Neutrophil Photoconversion

Photoconversion of neutrophils at the liver area were performed in anesthetized 13 dpf Tg(mpx:dendra) larvae fed with ND or HCD and mounted in a zWEDGI. For such, a 405-nm laser was focused into an oval area surrounding the liver for 45 seconds with 70% power, 10.0 μ/pixel. After photoconversion, tailfin amputation was performed and time-lapse movies from the wound area were performed from 1-6 hours post wounding acquiring one image every 15min, as described. Number of photoconverted neutrophils and percentage of photoconverted neutrophils at wound area at 1-, 2-, 4- and 6-hours post wounding were quantified manually using acquired time-lapse movies, as described.

### Drug treatments

Larvae were treated with metformin (Met) and pentoxifylline (PTX) as described previously^25^. Briefly, we dissolved metformin (Enzo Life Sciences) in E3 without methylene blue at a final concentration of 50 μM. PTX was first reconstituted in dimethyl sulfoxide and later diluted 1000× in E3 without methylene blue for a final concentration of 50 μM. Larvae were treated with these drugs for 72 hours, from 10-12 days post fertilization. Feedings were performed normally with cholesterol-enriched diet during drug treatments. Drugs were freshly prepared and replaced daily. At 12 dpf, drugs were washed and replaced with fresh E3 without methylene blue 24 hours prior to tailfin amputation to avoid direct effect of neutrophil recruitment during wounding assay. At 13 dpf, tailfin amputation was performed in larvae for all treatments. They were fixed at 4 hpw and imaged as described. Automatic quantification of neutrophils at wound area, systemic chronic inflammation (SCI), and neutrophil CHT depletion scorings were performed as described.

### Statistical analysis

All experiments were replicated independently two to three times (N) with multiple samples in each replicate (n). Least Squared Means analysis in R (www.r-project.org) ^21^ was performed on pooled replicate experiments, using Tukey method when comparing more than two treatments. For analysis of systemic chronic inflammation (SCI) and neutrophil CHT depletion we used Chi-Square test on GraphPad Prism version 8. Graphical representations were done in GraphPad Prism version 8. Statistical tests, p values, and n numbers used are given in each figure legends.

## Results

### NASH zebrafish larvae display systemic chronic inflammation

Inflammatory response triggered by fat accumulation and associated lipotoxicity is a major contributor for progression from simple steatosis to NASH REF. Increased number of circulating myeloid cells, particularly neutrophils, have been reported in NAFLD/NASH^10^.Using a diet induced NASH zebrafish model that exhibits liver inflammation ^14^ and metabolic syndrome^26^ after exposure to a high cholesterol diet (HCD) for 8 days (Suppl Fig. 1A), we first evaluated whether NASH larvae have increased number of myeloid cells. For such, we performed whole-larvae non-invasive confocal imaging at 13 days-post-fertilization (dpf) of double transgenic larvae expressing fluorescent nuclear markers in macrophages and in neutrophils [Tg(mpeg1:H2B-GFP; lyzC:H2B-mCherry)] fed with normal or HCD. We observed that NASH larvae fed with HCD have a significant increase in the total number of neutrophils and macrophages, 29% and 60% respectively, when compared to control larvae that were fed with the normal diet (Fig.1 A-C). Importantly, short exposure to the HCD did not have any impact on larvae development when compared with normal diet. The NASH and control larvae had similar width and length amongst the two groups (Suppl. Fig. 1). Nevertheless, larvae fed with HCD revealed mostly a severe systemic chronic inflammation (SCI) phenotype with enormous infiltration of neutrophils and macrophages into different tissues and organs (Fig. 1A and D; Suppl. Fig. 2). This was shown by automatic quantification of the number of these cells at different regions of the zebrafish larvae, particularly in the liver/posterior intestine, gastrointestinal track/heart and dorsal muscle/skin (Suppl. Fig. 3). We also observed a cytokine imbalance with several inflammatory mediators displaying altered gene expression levels in NASH conditions (data not shown). Interestingly, analysis of the caudal hematopoietic tissue (CHT) region, the main hematopoietic niche at this developmental stage, show drastic neutrophil depletion in NASH larvae fed with HCD (Fig 1A and E; Suppl. Fig.3B). We further decided to evaluate, how many days we needed to expose larvae to the HCD to induce SCI. Collecting larvae from 1 to 7 days of feeding (corresponding to 6 days-post-fertilization to 12 days-post-fertilization) we found that SCI starts to be observed at 4 days of feeding with severe SCI becoming the main phenotype observed by 7 days (Suppl. Fig. 4), these observations matched when we also start to observe severe steatosis and infiltration of neutrophils to liver area ^14^. Collectively, these findings indicate NASH is associated with increased number of myeloid cells and systemic chronic inflammation in zebrafish larvae.

**Figure 1:**
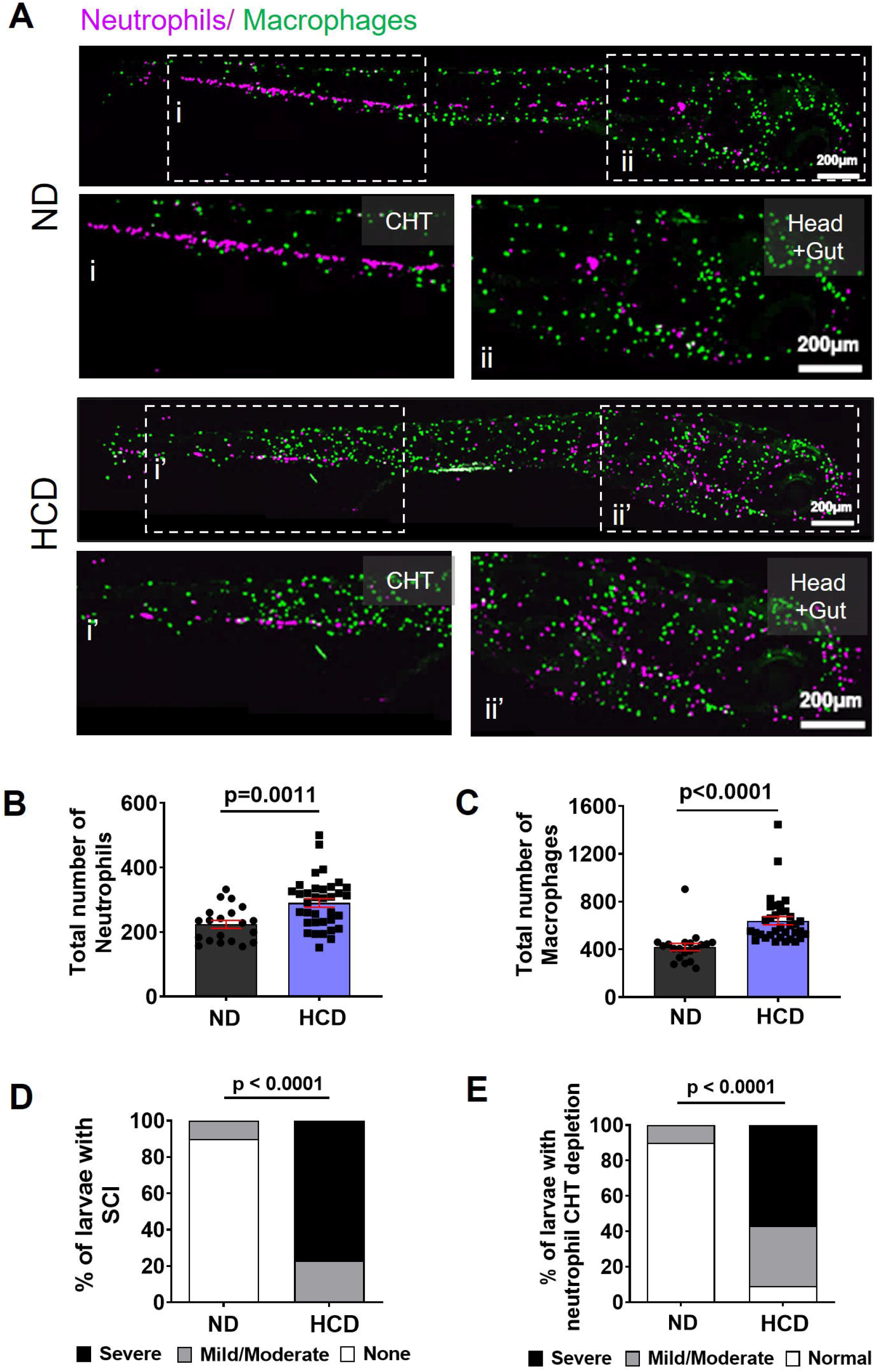
NASH larvae have enhanced myelopoiesis and systemic chronic inflammation. **(A)** Representative maximum intensity projections of 13 dpf larvae Tg(mpeg:H2B-GFP/lyzC:H2B-mCherry) fed with normal diet (ND) and high cholesterol diet (HCD); (i and i’) Higher magnification of the caudal hematopoietic tissue (CHT) dotted area. (ii and ii’) Higher magnification of the head and gut dotted area. **(B-C)** Quantification of total number of neutrophils (B) and macrophages (C) in whole-larvae (ND n=20, HCD n=35). **(D)** Chi-square graphs showing percentage of larvae with Systemic Chronic Inflammation (SCI) (ND n=20, HCD n=35). **(E)** Chi-square graphs showing percentage of larvae with neutrophil CHT depletion (ND n=20, HCD n=35). Data are from at least three independent experimental replicates. EM-Means analysis in R, was performed in total number of neutrophil and macrophage quantifications (B and C) and Chi-square test was used to analyze SCI and Neutrophil CHT depletion scorings (D and E). Scatter plots with bars shown mean ±SEM, each dot represents one larva, significant p values are shown in each graph. Scale bar = 200 µm.

### Proliferation of multipotent progenitors is enhanced in NASH

The consumption of Western-type diets trigger myelopoiesis and transcriptional reprogramming of myeloid precursor cells ^27^. Next, we decided to address if the increased number of neutrophils and macrophages found in NASH larvae were associated with increased proliferation at the caudal hematopoietic tissue (CHT) and the kidney, the hematopoietic niches where myelopoiesis occurs at this developmental stage^28^. To do so we incubated zebrafish larvae with EDU and measured the number of cells proliferating in the CHT, kidney, and thymus (Fig. 2). We found an increase in total proliferation in larvae fed with HCD compared to normal diet (Fig. 2 A and B). As expected, we also observed an increase in the number of proliferating cells at CHT and kidney, but not at the thymus (Fig. 2 A and C). In addition, we also observed an increased number of hematopoietic stem cells in larvae fed with the HCD, which was assessed by automatic quantification CD41+ cells in whole larvae (Suppl. Fig. 5). This data suggests that NASH larvae have increased myelopoiesis, explaining the increased number of myeloid cells observed.

**Figure 2:**
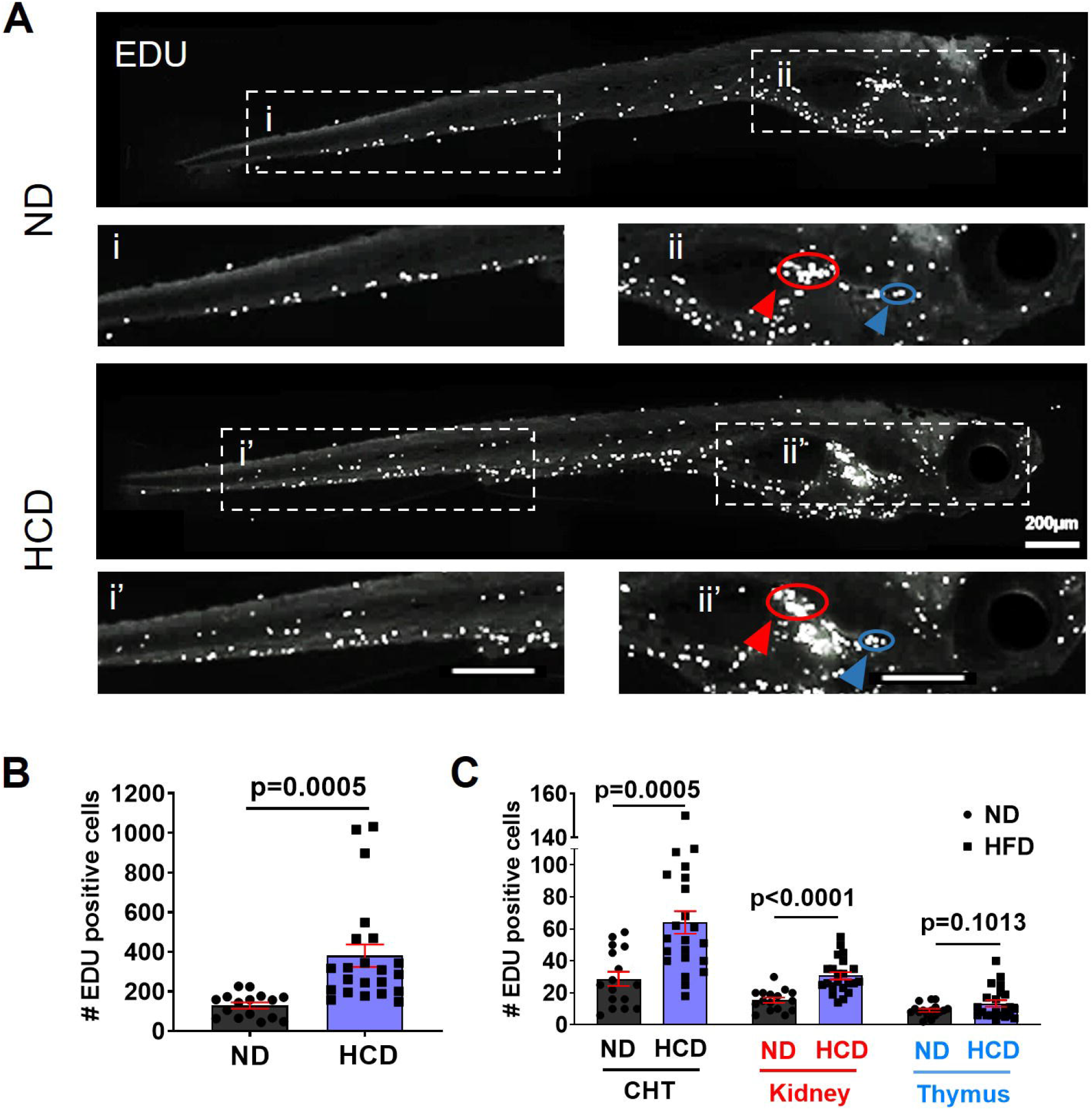
Proliferation in hematopoietic tissues is increased in NASH larvae. **(A)** Representative maximum intensity projections of 13 dpf larvae fed with normal diet (ND) and high cholesterol diet (HCD) incubated with EDU. **(B)** Quantification of number of EDU positive cells in whole larvae (ND n=16, HCD n=22). **(C)** Quantification of number of EDU positive cells at different hematopoietic tissues, Caudal Hematopoietic Tissue (CHT), Kidney and Thymus (ND n=16, HCD n=22). All data plotted are from at least two independent experimental replicates. EM-Means analysis in R was performed in all data. Scatter plots with bars shown mean ±SEM, each dot represents one larva, significant p values are shown in each graph. Scale bar= 200μm.

### Neutrophil recruitment to a tailfin injury is exacerbated in NASH

Multiple studies have shown that metabolic syndrome and metainflammation trigger a hyperactive response of myeloid cells ^27, 29, 30^. To test whether NASH zebrafish larvae display a hyperactive neutrophil response, we used a well-established zebrafish tailfin injury model^16, 19^ to investigate how NASH impacts neutrophilic inflammation. Using tailfin injury as a “secondary” inflammatory trigger, we observed an exacerbated neutrophil response to the tailfin injury in larvae fed with HCD compared to normal diet with 2.8 times more neutrophils recruited at the peak of recruitment (Fig. 3; Supplemental Movie 1). This exacerbated neutrophil recruitment in NASH larvae was not accompanied by macrophage infiltration, with macrophages being recruited to tailfin injury at same level as in control larvae (Supplemental Movie 1). Moreover, analysis of neutrophil tracking for 8 hours, showed that neutrophils migrate at a higher speed at the wound and vicinity in NASH larvae (Fig. 4 A-B). Interestingly, at this developmental stage a vast majority of the neutrophils that arrive to the tailfin area use vessels, particularly in NASH larvae. We could observe this in the time-lapse movies, by detecting the accumulation of neutrophils at the artery-vessel loop (Fig. 3; Supplemental Movie 1). Collectively, our findings show that cholesterol surplus and presence of NASH promotes an exacerbated neutrophil response to tissue injury, supporting the idea that western type diets and associated pathologies promote alterations on neutrophil biology.

**Figure 3:**
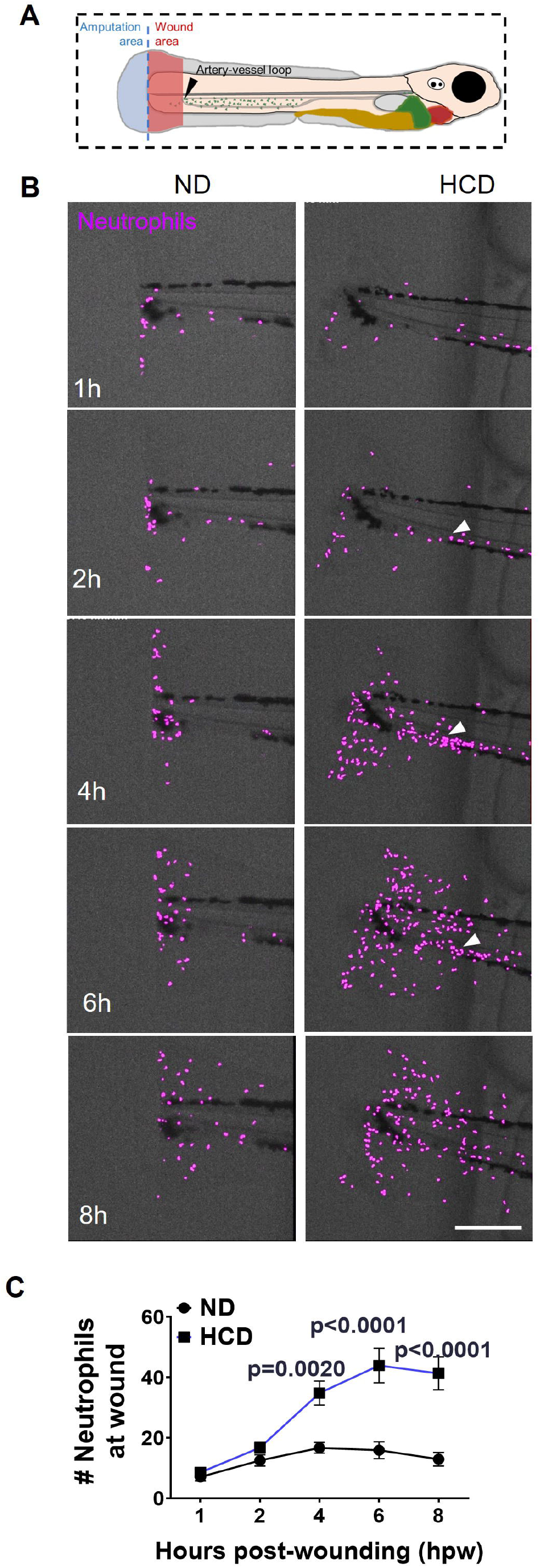
Neutrophil response to tissue injury is enhanced in NASH larvae. **(A)** Representative maximum intensity projections of 13 dpf larvae Tg(lyzC:H2B-mCherry) fed with normal diet (ND) and high cholesterol diet (HCD). Images were extracted from time-lapse movies **(B)** Quantification of number of neutrophils recruited to wound at 1-, 2-, 4-, 6- and 8- hours-post wounding [(hpw); (ND n=16, HCD n=17)]. Data are from at least three independent experimental replicates. Two-way-ANOVA analysis with Bonferroni’s multiple comparisons test. Solid line shows mean ±SEM, significant p values are shown in each timepoint. Each dashed line in graph represents neutrophil recruitment to wound area in one larva followed from 1-8hpw. Scale bar= 200μm.

**Figure 4:**
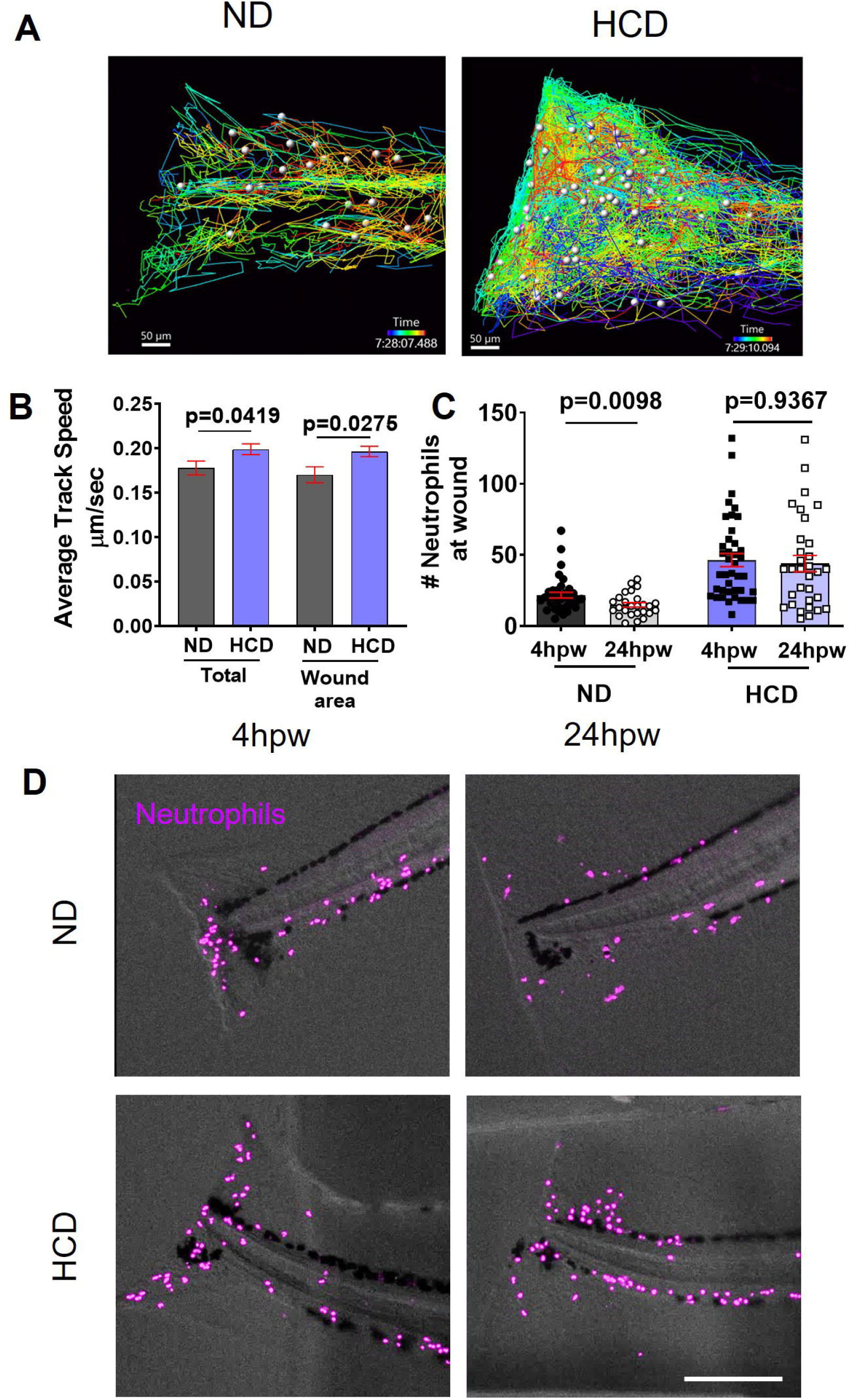
Resolution of inflammation is delayed in NASH larvae. **(A)** Representative maximum intensity projections of tailfin wounds at 4 and 24 hours-post-wounding (hpw) of 13 dpf larvae Tg(lyzC:H2B-mCherry) fed with normal diet (ND) and high cholesterol diet (HCD). **(B)** Quantification of number of neutrophils recruited to wound at 4 and 24 hours post wounding [(hpw); (ND 4hpw n=36, ND 24hpw n=26, HCD 4hpw n=40, HCD 24hpw n=32)]. Data are from at least three independent experimental replicates. EM-Means analysis in R was performed. Scatter plots with bars shown mean ±SEM, each dot represents one larva, significant p values are shown in each graph. Scale bar= 200μm.

### Resolution of inflammation in tissue injury is impaired in NASH

Resolution of inflammation is a critical phase in the inflammatory response, where neutrophil clearance from the injured area needs to occur so that the tissue repair machinery can be fully activated. Once resolution of inflammation is dysregulated, progression from acute to chronic inflammation occurs and tissue damage and disease results^31^. The kinetics of the recruitment curve from time-lapse microscopy were different in NASH larvae compared to control suggesting that neutrophil clearance from the wound is delayed and therefore possibly impacting resolution of inflammation (Fig. 3B). To test whether resolution phase is affected in NASH, we quantified the number of neutrophils at the tailfin wounded area at 4 hpw, the peak of neutrophil recruitment, and later at 24 hpw, when resolution phase is expected to have started in larvae fed with HCD versus normal diet ^16, 32-34^. As expected, we observed a reduction in the number of neutrophils at the wound from 4 to 24 hpw in control larvae fed with normal diet (Fig. 4 C-D). However, in NASH larvae fed with HCD the number of neutrophils at the injury site were almost the same at 4 and 24 hpw (Fig. 4 C-D). Collectively, these findings show that NASH impairs resolution of inflammation in tissue injury.

### Neutrophils from NASH larvae have increased ROS production at injury sites

Next, we decided to evaluate if exacerbated neutrophil chemotaxis and impaired resolution of inflammation were associated with increased ROS production in neutrophils at the tissue injury site. To do so we stained larvae expressing a neutrophil cytoplasmic marker, Tg(mpx:mCherry), with CellROX Deep Green to label intracellular H_2_O_2_, and performed colocalization analysis on confocal microscopy images of the amputated tailfin (Fig. 5). We observed that larvae fed with HCD have a significantly higher number of neutrophils at the wound site that generate ROS at 4 hpw, as shown by the quantification of the number of colocalized voxels between neutrophil signals (mCherry) and CellROX (Deep Green) (Fig. 5 A and B). In addition, we also found that neutrophils from larvae fed with HCD generate higher amount of H_2_O_2_, as shown by quantification of the percentage of mCherry/Deep Green signal that colocalizes (Fig. 5 C). Collectively, these results indicate that NASH enhances the ability of neutrophils to produce a higher amount of ROS at inflammatory sites, suggesting that NASH induces neutrophil priming.

**Figure 5:**
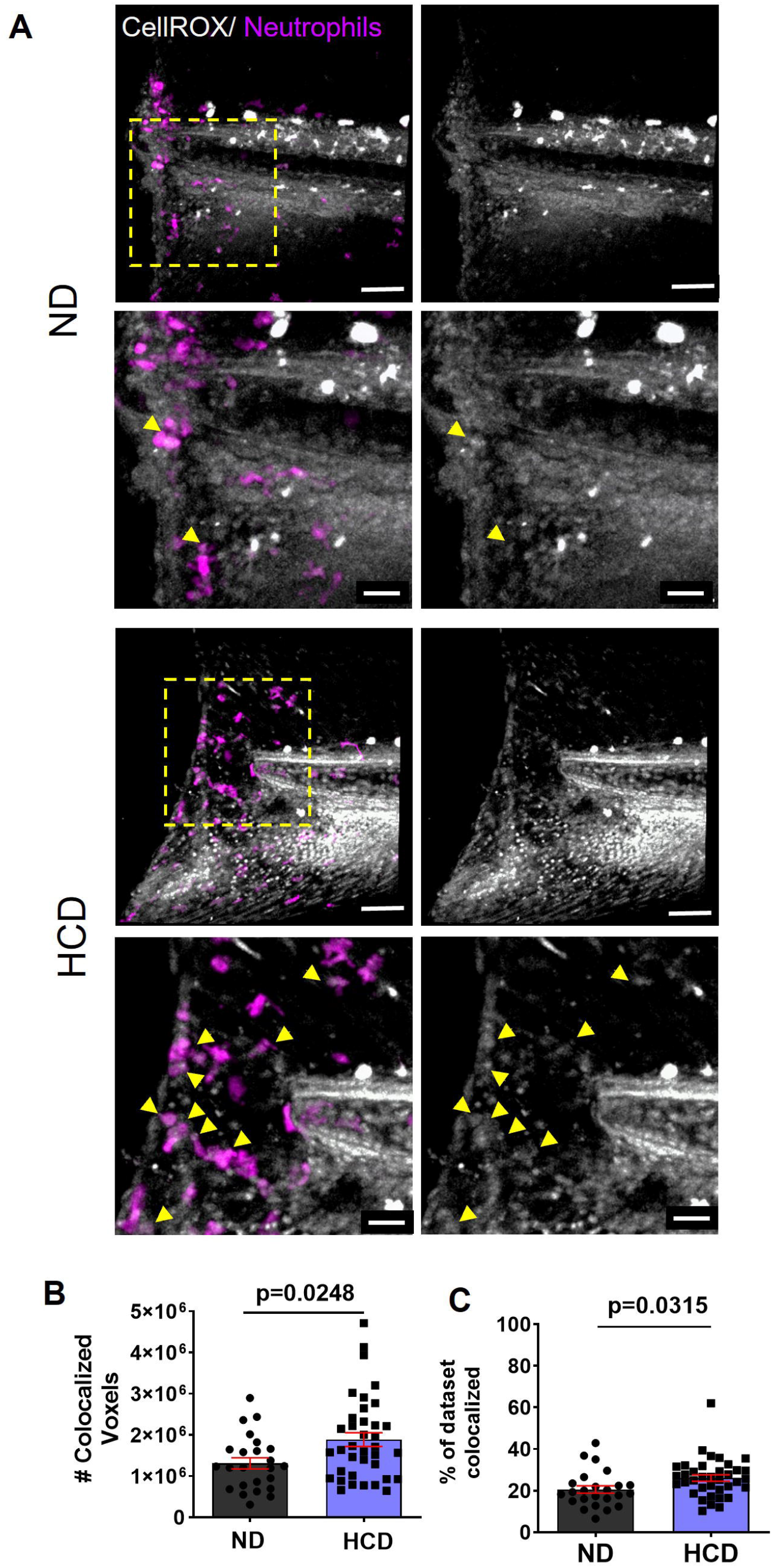
NASH enhances neutrophil ROS production at injury sites. **(A)** Representative maximum intensity projections of tailfin wounds at 4hours-post-wounding (hpw) of 13 dpf Tg(mpx:mCherry) larvae fed with fed with normal diet (ND) and high cholesterol diet (HCD) and incubated with CellROX. Scale bar= 50 μm. Higher magnification of dotted area at tailfin wound. Scale bar = 20 μm. **(B)** Quantification of number of colocalized voxels. (C) Quantification of the % of dataset colocalized (ND n=24, HCD n=37). Data are from at least three independent experimental replicates. EM-Means analysis in R was performed. Scatter plots with bars shown mean ±SEM, each dot represents one larva, significant p values are shown in each graph.

### Neutrophils reverse migrate from liver area to tailfin injury

Clearance of neutrophils from inflamed areas is achieved by multiple fates, such as apoptosis, phagocytosis, or reverse migration ^35^. Neutrophil reverse migration is a conserved mechanism among human, murine and zebrafish. In a mice model of sterile liver injury, a subpopulation of neutrophils leaves the injury site by reverse migration and re-enter the circulation REF. Next, we decided to test whether neutrophils infiltrated at the NASH liver ^14, 15^ can reverse migrate and respond to the “*secondary”* inflammatory stimulus, the tailfin injury. For such, we performed photoconversion of neutrophils at the liver area (Suppl. Fig. 5) making use of a transgenic line [Tg(mpx:dendra)] with neutrophils expressing the photoconvertible protein Dendra. Immediately after photoconversion, a tail fin amputation was performed followed by non-invasive time-lapse confocal microscopy imaging from 1-6 hpw. We observed that larvae exposed to HCD had a higher number and higher percentage of photoconverted neutrophils at the wound from 1-6 hpw (Fig. 6). This data suggests that neutrophils from NASH liver can undergo reverse migration and massively respond to a *“secondary”* inflammatory stimulus. Such effect was also observed in our whole-animal time-lapse movies (Supplemental Movie 3).

**Figure 6:**
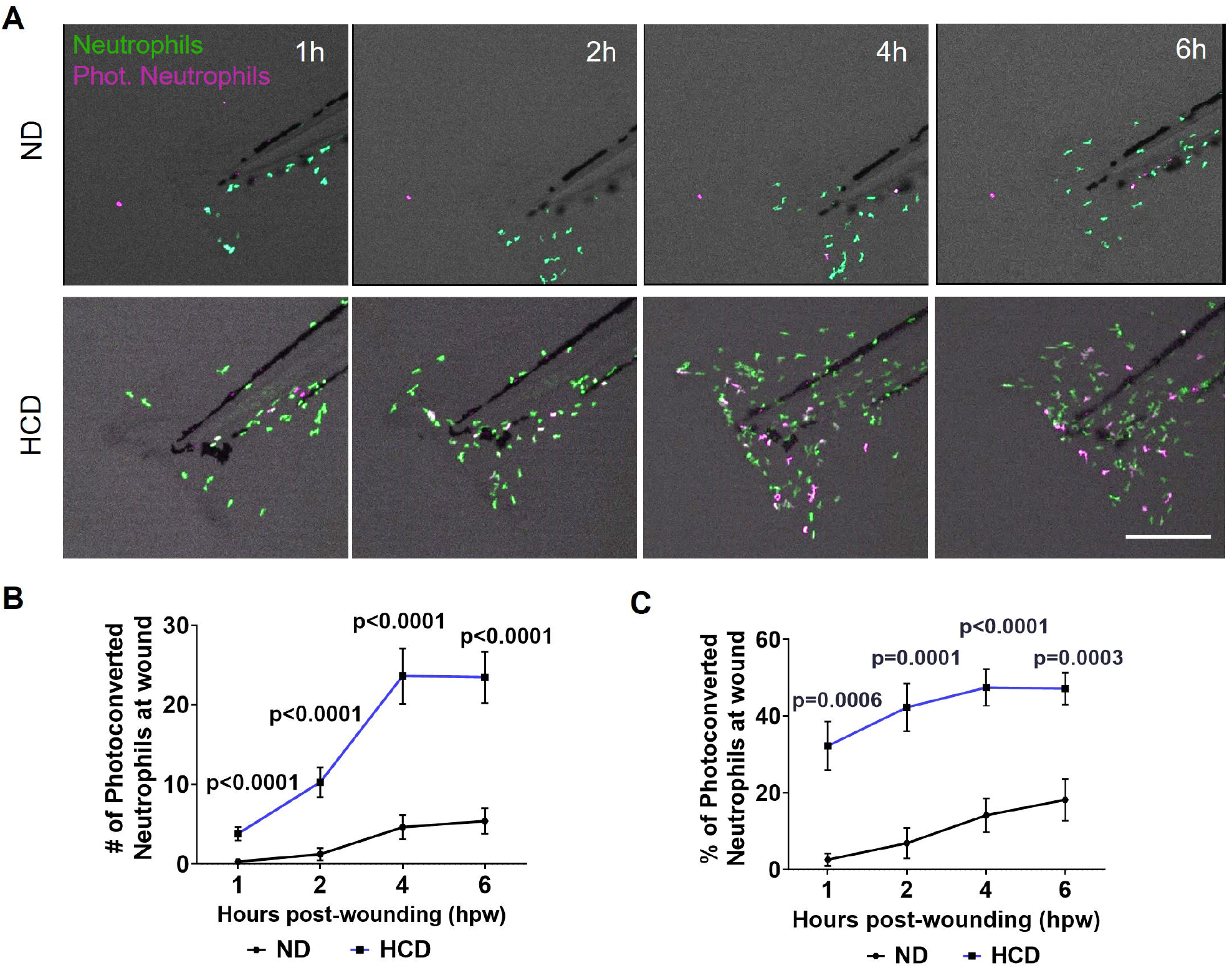
Neutrophils undergo reverse migration from inflamed NASH-liver to tailfin injury. **(A)** Representative maximum intensity projections of 13 dpf larvae Tg(mpx:Dendra) fed with normal diet (ND) and high cholesterol diet (HCD). Images extracted from time-lapse movies. **(B)** Quantification of number of photoconverted neutrophils recruited to wound area at 1-, 2-, 4- and 6-hours post wounding [(hpw); (ND n=15, HCD n=20)]. **(C)** Quantification of percentage of photo-converted neutrophils recruited to wound area at 1-, 2-, 4- and 6-hpw (ND n=15, HCD n=20). Data are from at least three independent experimental replicates. Two-way-ANOVA analysis with Bonferroni’s multiple comparisons test. Solid line shows mean ±SEM, significant p values are shown in each timepoint. Each dashed line in graph represents neutrophil recruitment to wound area in one larva followed from 1-6hpw. Scale bar= 200μm.

### Neutrophil exacerbated response to tissue injury is alleviated by NASH pharmacological intervention

In our previous studies, metformin treatment reduced steatosis, and overall inflammation at the liver that could be observed by reduced neutrophil infiltration ^14^. In addition, tumor necrosis factor-alpha (TNFα) has been reported as a main inflammatory molecule upregulated in NASH, as we have shown previously shown in our NASH model ^14^. Inhibition of TNFα secretion with pentoxifylline ^36^, was found to be effective on improving liver function and histological changes in patients with NASH To test whether pharmacological treatment of NASH with metformin and pentoxifylline could revert the SCI as well the exacerbated neutrophil response, we first let larvae to develop NASH and SCI by feeding larvae from 5 days post fertilization (dpf) to 9 dpf with HCD (Suppl. Fig. 2), followed by a 2-day treatment (10 dpf-12 dpf) with metformin or pentoxifylline to revert this effect. Drug treatments were removed and replaced by E3 at least 16h before any intervention in larvae to avoid direct effect on neutrophil recruitment to tailfin injury. We observed that both these drugs partially rescued diet-induced SCI and decreased the hyper-responsiveness of neutrophils to tailfin injury (Fig. 7), suggesting that indeed NASH and associated liver inflammation contributes to priming of neutrophils and that NASH-pharmacological intervention can alleviate the adverse effect on exacerbated neutrophilic inflammation.

**Figure 7:**
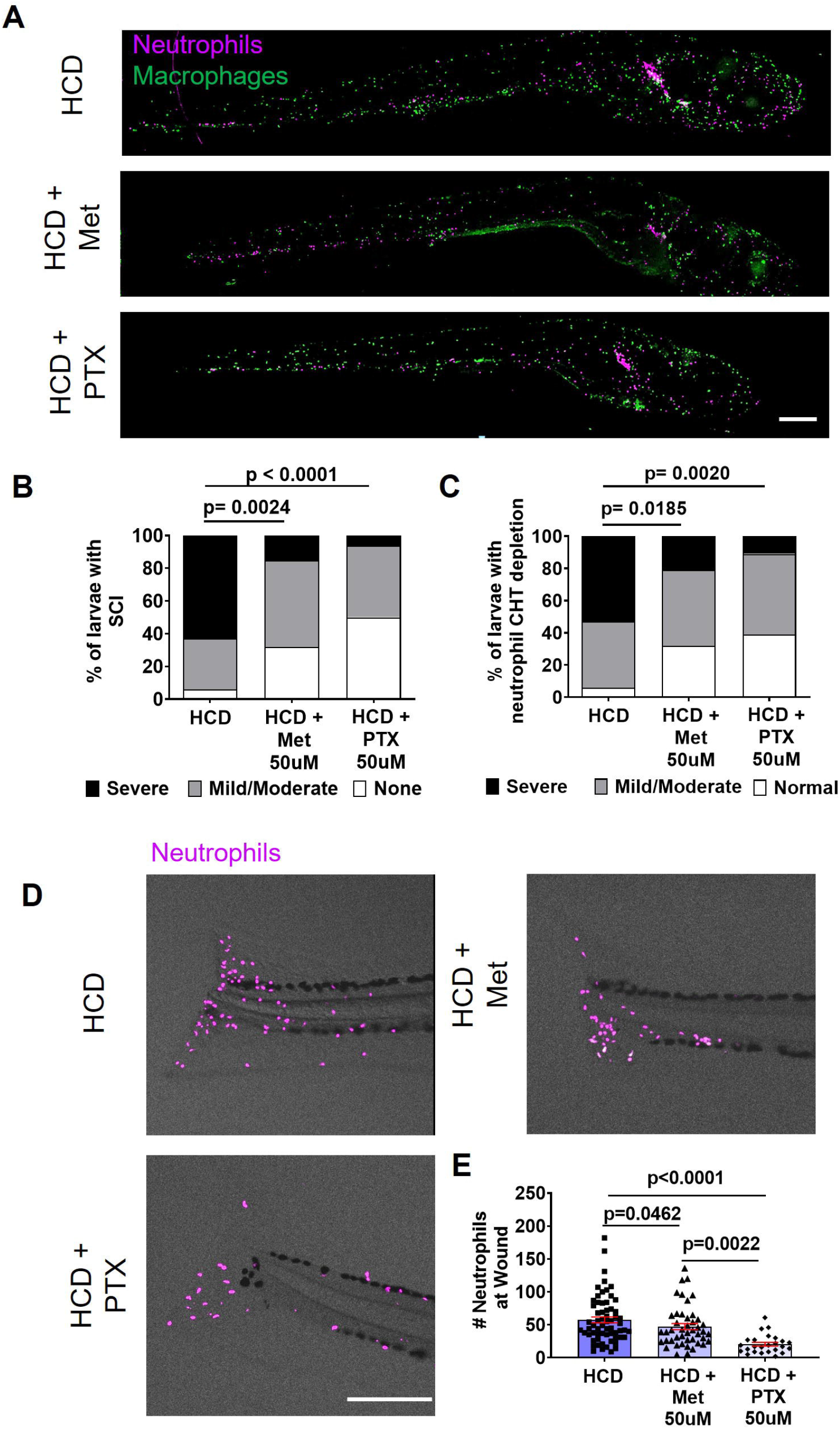
Pharmacological treatment with Metformin and Pentoxifylline alleviates NASH impact on neutrophilic inflammation. **(A)** Representative maximum intensity projections of 13 dpf larvae Tg(mpeg:H2B-GFP/lyzC:H2B-mCherry) fed with normal diet (ND) and high cholesterol diet (HCD) treated with DMSO, Metformin (Met) or Pentoxifylline (PTX). **(B)** Chi-square graphs showing percentage of larvae with Systemic Chronic Inflammation (SCI) (HCD n=32, HCD+Met n=19, HCD+PTX n=18). **(C)** Representative maximum intensity projections of tailfin wounds at 4 hours-post-wounding (hpw) of 13 dpf larvae Tg(lyzC:H2B-mCherry) fed with ND and HCD and treated with DMSO, Metformin (Met) or Pentoxifylline (PTX). **(D)** Quantification of number of neutrophils recruited to wound area at 4hpw (HCD n=63, HCD+Met n=45, HCD+PTX n=25). Data are from at least three independent experimental replicates. EM-Means analysis in R, was performed in quantification of number of neutrophils at wound (B) and Chi-square test was used to analyze SCI and Neutrophil CHT depletion scorings (D and E). Scatter plots with bars shown mean ±SEM, each dot represents one larva, significant p values are shown in each graph. Scale bar = 200 µm.

## Discussion

Nonalcoholic fatty liver disease (NAFLD), and its more aggressive inflammatory form, nonalcoholic steatohepatitis (NASH), are associated with metainflammation^37, 38^ and increased activation of neutrophils^39^. How exactly systemic neutrophilic inflammation is altered by NASH and associated metainflammation and what mechanisms sustain neutrophil hyperactive response are not fully understood. A major limitation in the field is the lack of vertebrate animal models of NASH amenable to whole-animal non-invasive live-imaging of immune cell recruitment and function that recapitulate the inherent cellular and molecular complexity of establishing inflammatory responses in the context of metabolic syndrome and metainflammation. The transparency and easiness to perform live imaging makes the zebrafish the only vertebrate system that allows the visualization and study of neutrophils and inflammatory response non-invasively in a whole-animal context^13^. Here making use of a diet induced-NASH zebrafish model we report that neutrophil response to tissue injury is drastically exacerbated with neutrophils from the liver undergoing massive reverse migration upon sensing a “secondary” inflammatory stimulus, functioning as a main source of primed neutrophils that fuels neutrophilic inflammatory response to injury sites. We also probe that pharmacological treatment of NASH reverted the observed exacerbated neutrophil response to tissue injury. Our work shows that zebrafish models provide the perfect platform to study the pathophysiological mechanisms involved on how diet, NASH and associated metainflammation impact neutrophils and inflammatory response. Overall, our findings support the idea that under NASH conditions, the liver serves as a source of primed neutrophils that reverse migrate and respond massively to “secondary” inflammatory triggers, identifying a potential therapeutic target to reduce the adverse complications observed in patients with NASH and associated comorbidities due to hyperactive neutrophil response.

The liver is a vital organ with high regenerative capacity that plays more than 500 functions, with a central role in metabolic activities, nutrient storage, detoxification^40^. The liver is also the stage for complex immunological actions; being exposed to dietary and commensal bacterial products from the gut with inflammatory potential that routinely challenge the diverse population of resident immune cells. Interestingly, the liver not just facilitates the removal and degradation of immunogenic molecules from the gut^40^, but it is also target of inflammatory macromolecules from the brain via a drainage mechanism that results in macrophage and neutrophil infiltration^41^. To exert its functions, the liver tolerates those challenges but at same time triggers homeostatic inflammation, a process that is continuously being activated and resolved to support tissue regeneration and preserve tissue and organ homeostasis^40^. Once liver homeostatic inflammation is dysregulated, pathology and organ damage occur^40^.The consumption of high cholesterol diet can be one of the factors that disrupts homeostatic inflammation; indeed, cholesterol surplus leads to polarization of Kupfer Cells (the resident macrophage population) into a M4-like phenotype that promotes recruitment of neutrophils^11^. Without the possibility to shut down and resolve inflammation, chronic liver inflammation is established, which gives raise to NASH that eventually develops to fibrosis and cancer. We and others have shown that zebrafish larvae develop NASH after exposure to high cholesterol diet (HCD) ^14, 15, 26, 42^. Systemic inflammation and overall immune imbalances, both at cellular and cytokine levels, are associated with NASH and other metabolic diseases^4, 43^. We found that NASH larvae develop a chronic systemic low-grade of inflammation characterized by infiltration of neutrophils and macrophages to different tissues and an imbalance in the expression of several pro-inflammatory cytokines. In addition, we observed enhanced proliferation at hematopoietic niches and number of hematopoietic stem cells, which explained the increased number of neutrophils and macrophages in NASH larvae. NASH can contribute to establishment of such chronic low-grade systemic inflammation, which ultimately triggers myelopoiesis, through the systemic release of several markers of inflammation, oxidative stress, and of procoagulant factors. Our observations phenotypically recapitulate the systemic impact of NASH found in humans and murine models^43, 44^, providing a basis for the use of the NASH zebrafish model on evaluating the impact of this pathology on neutrophil response to “secondary” inflammatory triggers.

One of the best characterized models of live neutrophil recruitment is the zebrafish tailfin injury model, which triggers a leukocyte immune response that precisely mimics the kinetics observed in mammalian acute inflammatory responses ^13, 16, 19, 32, 45-55^. We found that neutrophil response to tissue injury is exacerbated in presence of NASH and associated metainflammation, with a substantial number of neutrophils being recruited (2.8 times more) after exposure to HCD. Similarly, exacerbated neuroinflammation and intracerebral hemorrhage injury was previously observed in a mouse model of NAFLD^56^. Moreover, recruited neutrophils to tissue injury migrate at a higher speed in NASH larvae. It has been reported in multiple models that cholesterol surplus increases adherence and decreases rolling velocity of neutrophils. As far as we know, this is the first time a study reports a faster speed of neutrophils migrating in an injured tissue and vicinity under NASH conditions. Increased neutrophil speed at injury sites could seriously impact the local inflammatory response interfering with neutrophil recognition of tissue injury site. The increased neutrophil speed observed in NASH and associated metainflammation, could be a result of low production of chemokines responsible for slowing-down neutrophils at injury sites, as we and *Sarris et al* have shown previously with CXCL8 ^16, 57^, due to cell exhaustion or tolerance. Another possible explanation is that neutrophils from NASH larvae might have been through mechanisms of receptor desensitization, internalization, or degradation^58^ triggered by the chronic inflammation found in this condition. Such alterations of chemoattractant receptors would hamper neutrophils from recognizing the high levels of chemoattractants such as CXCL8 that indicate the localization of injury sites and allow neutrophils to engage in local inflammatory response.

Neutrophils can modify their functional responses after being exposed to multiple factors, through the process named neutrophil priming^59^. NASH liver is a chronic inflammatory microenvironment that can lead to neutrophil priming. In NASH and hyperlipidemic patients, peripheral polymorphonuclear leukocytes (PMNL) are primed and able to produce increased amounts of ROS^60, 61^. We observed in NASH larvae that neutrophils recruited to the tissue injury produce higher amounts of reactive oxygen species (ROS) at the injury site. Increased ROS production can cause a progressive oxidative damage, sustain inflammation, and delay resolution. Therefore, the increased ROS production levels observed in NASH neutrophils at the injury site could be one contributing factor for the delayed resolution that we observed. In addition, we found that nearly 50% of the neutrophils recruited to the tailfin injury have NASH liver as origin. Interestingly, pharmacological treatment of NASH with Metformin and Pentoxifylline reverted the systemic chronic inflammation and the exacerbated neutrophil recruitment to the tissue injury. Overall, these findings make us speculate that NASH liver is an active source of primed neutrophils that massively reverse migrate towards “secondary” inflammatory stimulus such as tissue injury; therefore, the neutrophil pool at NASH liver might be a potential target to reduce the adverse effects caused by dysfunctional and hyperactive neutrophil response observed in patients with NASH and associated comorbidities that often lead to disability and death.

Metabolic syndrome and associated inflammation is a complex interplay of signals among different tissues and organs that could all be contributing to the exacerbated neutrophilic inflammation observed in NASH larvae. Our study did not allow us to separate the effect of NASH from systemic chronic inflammation. It is possible that under the systemic chronic inflammation conditions, increased production of ROS and other proinflammatory signals from epithelia cells or immune resident cells like macrophages, might be contributing to the exacerbated neutrophil response found in NASH larvae. In our study, we observed infiltration of neutrophils to multiple tissues and organs and a drastic neutrophil depletion from hematopoietic tissues. We found that just about 50% of neutrophils at tailfin injury have reverse migrated from the liver, it is plausible to consider that other tissues and organs might also contribute to the impaired neutrophil response in NASH via similar mechanism. Additionally, it is unclear at what extension the cholesterol diet is impacting neutrophil biology directly. Lipids accumulate in leukocytes of rats fed with different atherogenic diets^62^. Neutrophil *ex vivo* treatment with for example cholesterol, low density lipoprotein or oxysterols support a direct role of cholesterol surplus on neutrophil chemotaxis, adhesion, function and fate^63-65^. In this study we did not explore such effect to determine at what extent the changes in neutrophil recruitment are cell autonomous or not. Finally, the metabolic and epigenetic rewiring of myeloid progenitor cells by diet drives trained immunity and sustains hyper responsiveness of the innate immune system^27^. Furthermore, different tissues and inflammation induce proteomic, transcriptomic and epigenomic reprogramming of neutrophils ^66^ REFS. Therefore, another important question that this study raises is at what cellular stage (e.g., progenitor level, immature, or mature), where (e.g., hematopoietic tissues, liver or other tissues and organs) and how neutrophils are being altered in NASH. Future studies will need to be performed to specifically address these questions and the diet-induce NASH zebrafish model is a unique model to visualize and investigate the cellular and molecular mechanisms that drive neutrophil hyperactive response in NASH.

In summary, our data suggest that NASH exacerbates neutrophilic inflammation to tissue injury probably via neutrophil priming at the liver, which can further undergo reverse migration and respond to secondary inflammatory triggers. In the future, reverse migration of neutrophils from the liver might be an important mechanism to target to diminish neutrophil response, improve prognosis, and reduce disability and death in patients with NASH.

## Abbreviations used in this paper

NAFLD: Nonalcoholic fatty liver disease
NASH: nonalcoholic steatohepatitis
ROS: reactive oxygen species
NETs: Neutrophil Extracellular Traps
NETosis: Neutrophil Extracellular Traps formation
HCD: high-cholesterol diet
TNF: tumor necrosis factor
dpf: days post-fertilization
ND: normal diet
Met: Metformin
Ptx: Pentoxifylline

## Acknowledgments

We thank to Dr. Anna Huttenlocher for laboratorial and financial support while conducting experimental work in UW-Madison and Dr. Veronika Milkosvki for all the critical comments and support given during the development of this work.

