## Supplemental Figures for "Neutrophil reverse migration from liver fuels neutrophilic inflammation to tissue injury in Nonalcoholic Steatohepatitis"

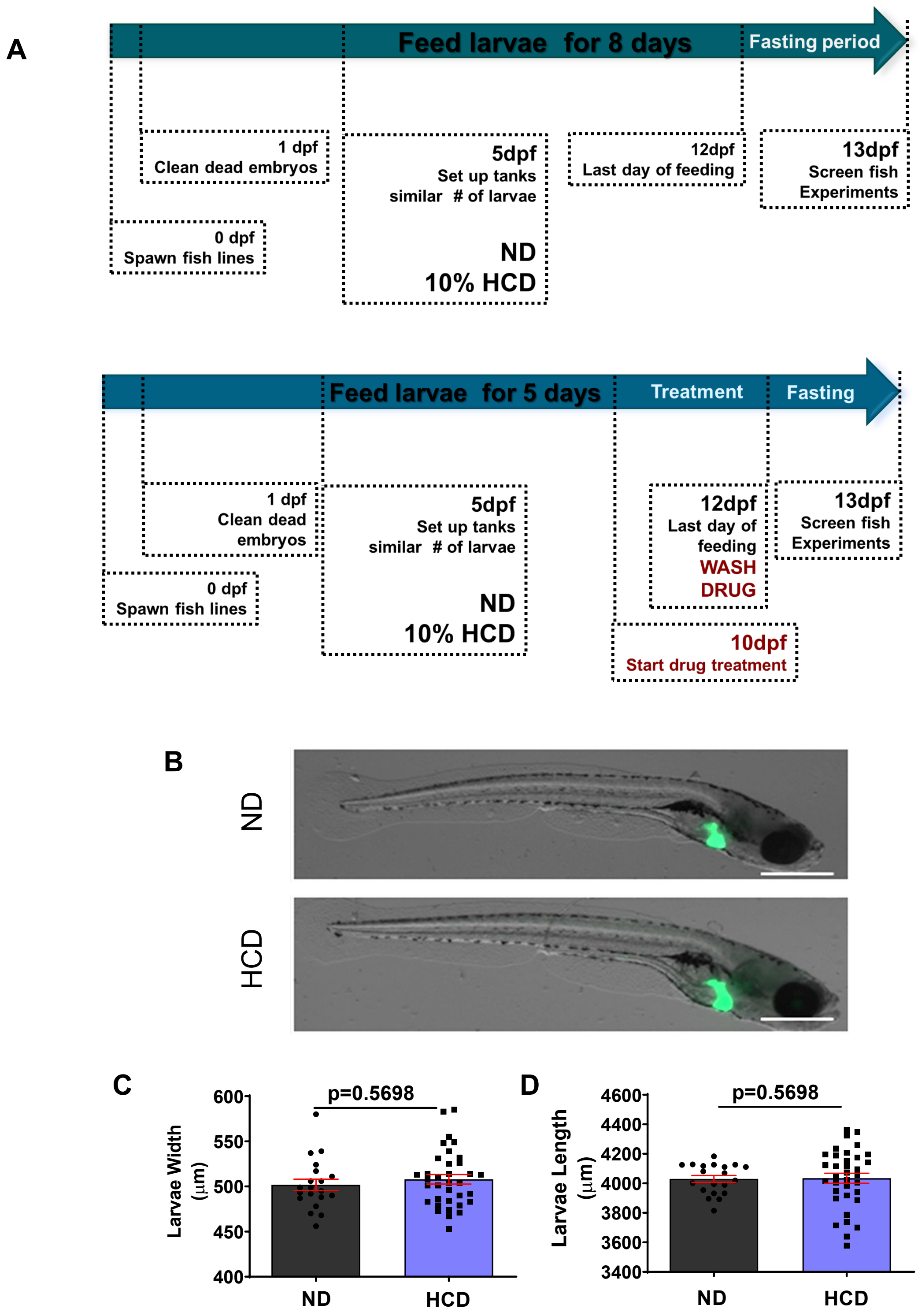


**Supplemental Figure 1: Short-term high cholesterol diet does not affect larvae development. (A)** Diagram of short-term high cholesterol diet (HCD). **(B)** Representative maximum intensity projections of 13 days-post fertilization (dpf) larvae fed with normal diet (ND) and high cholesterol diet (HCD). **(C-D)** Quantification of larvae width (C) and length (D). All data plotted comprise at least three independent experimental replicates. LS-Means analysis in R was performed. Scatter plots with bars shown mean ±SEM, each dot represents one larva, significant p values are shown in each graph, significant p values are shown in each graph. Scale bar= 500µm.

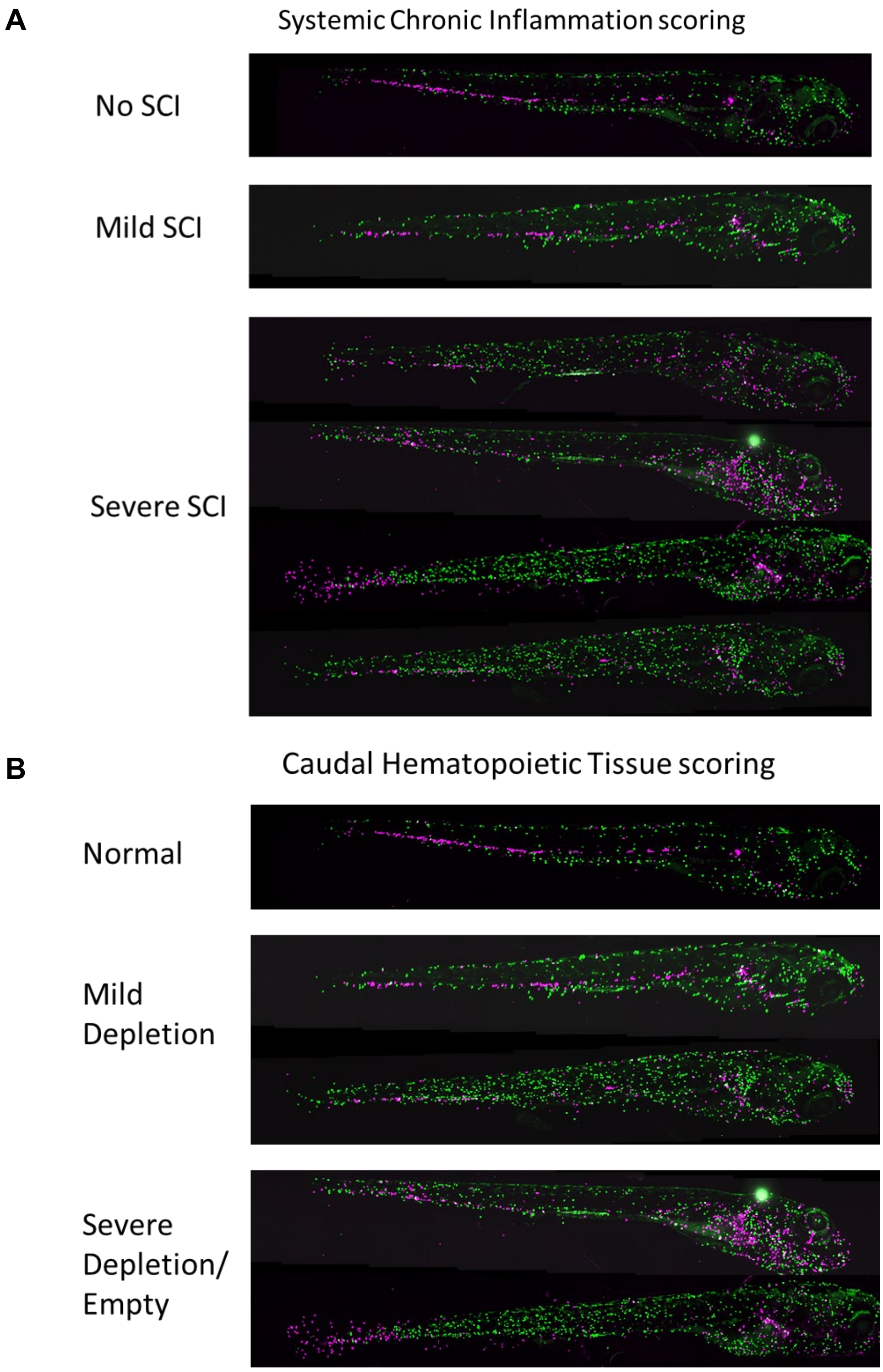


**<u>Suppl. Figure 2:</u>** Systemic Chronic Inflammation (SCI) and CHT neutrophil depletion scoring systems. (**A-B**) Representative maximum intensity projections of 13 days-post fertilization (dpf) transgenic larvae fed with normal diet (ND) and high cholesterol diet (HCD).

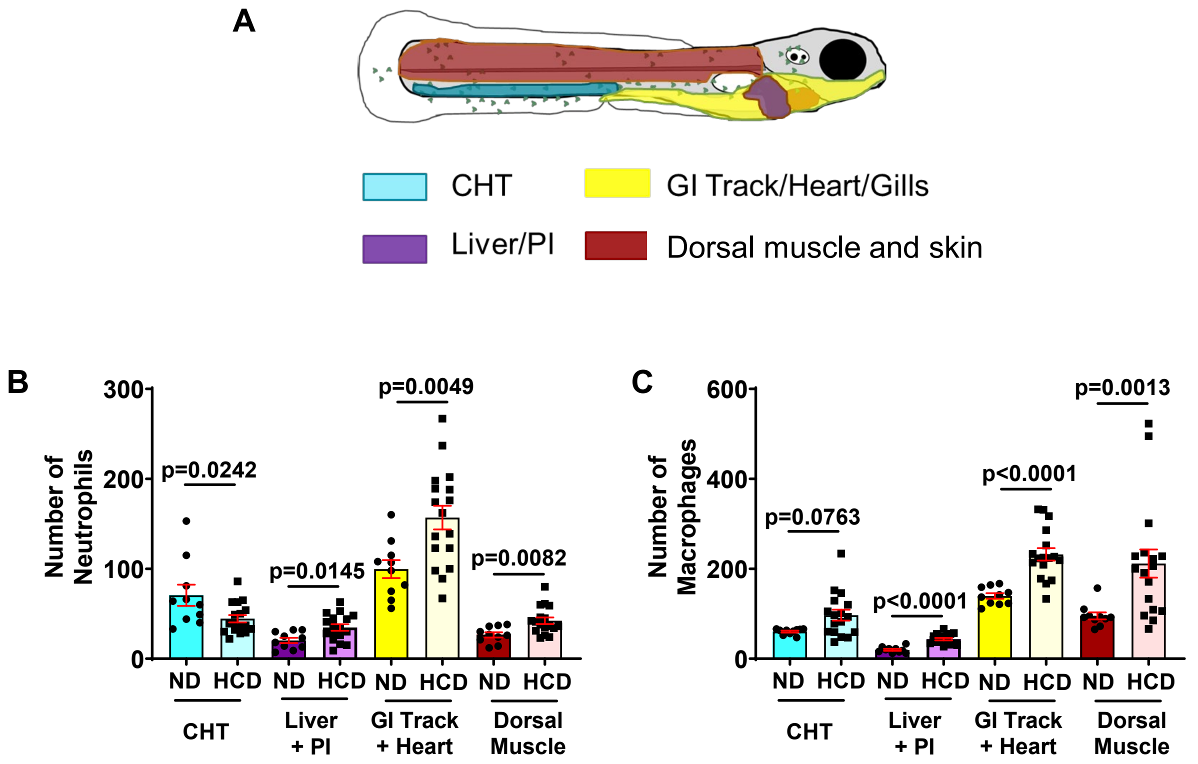


<u>**Suppl. Figure 3:**</u> **Short-term high cholesterol diet induces innate immune cell infiltration to tissue and organs. (A)** Diagram of different areas in the fish. **(B and C)** Quantification of number of neutrophils (B) and macrophages (C) present at different areas. Caudal Hematopoietic Tissue (CHT); liver and posterior intestine (PI); gastro-intestinal (GI) track, heart and gills; Dorsal Muscle. Analysis performed in Graph-Prism. t-test two-tailed with Mann Whitney post-test. Dot plots show mean ± SEM, p values are shown in each graph.

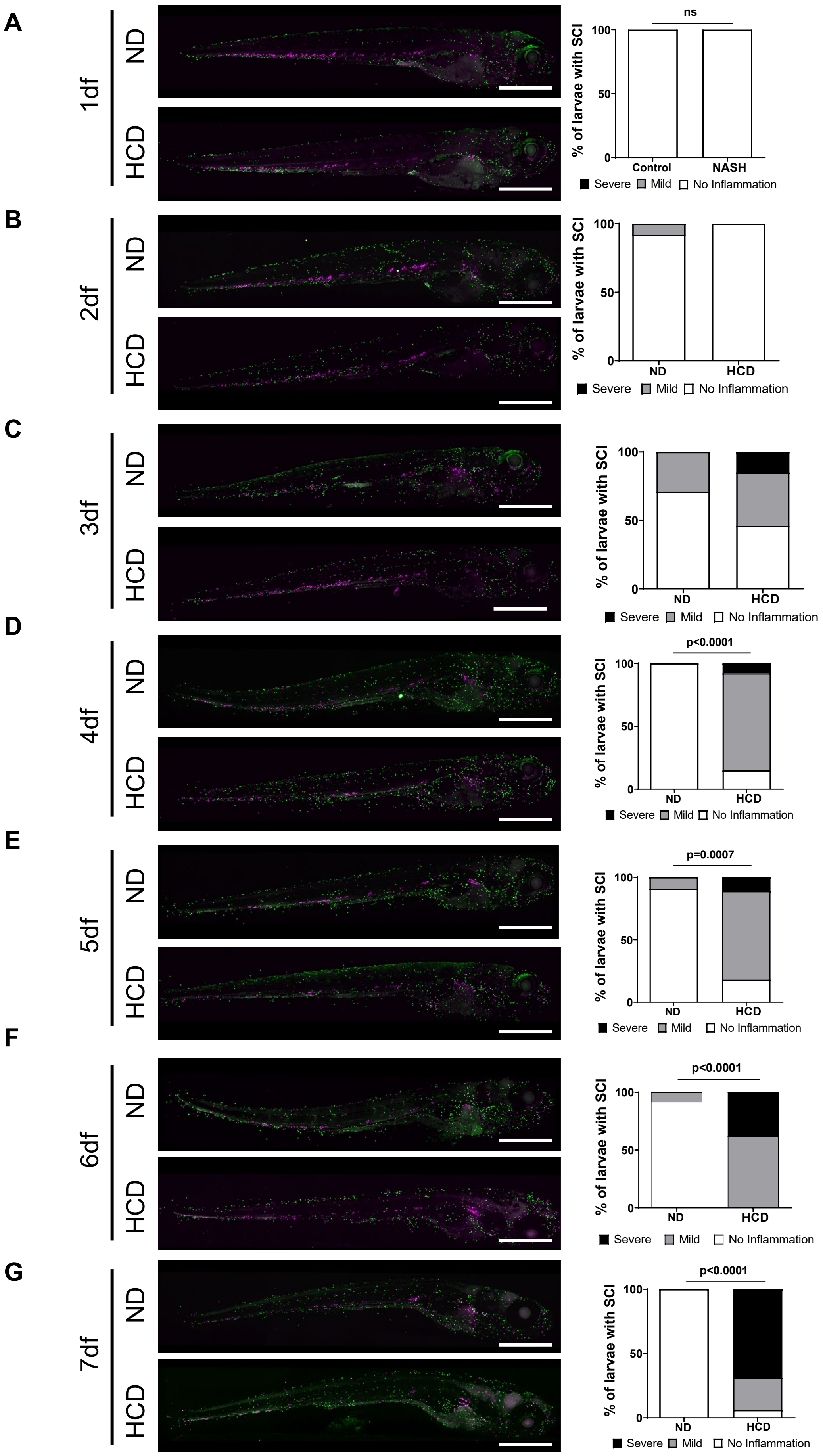
A
1df
ND
HCD
ns
% of larvae with SCI
100
50
0
Control
NASH
Severe Mild No Inflammation
B
2df
ND
HCD
% of larvae with SCI
ND
HCD
Severe Mild No Inflammation
C
3df
ND
HCD
% of larvae with SCI
Severe Mild No Inflammation
D
4df
p<0.0001
% of larvae with SCI
Severe Mild No Inflammation
E
5df
p=0.0007
% of larvae with SCI
Severe Mild No Inflammation
F
6df
p<0.0001
% of larvae with SCI
Severe Mild No Inflammation
G
7df
p<0.0001
% of larvae with SCI
Severe Mild No Inflammation

**Suppl. Figure 4:** Timeline of Systemic Chronic Inflammation (SCI). (**A-G**) Representative maximum intensity projections of 6-12 days-post fertilization (dpf) transgenic larvae fed with normal diet (ND) and high cholesterol diet (HCD) from 1-7 days of feeding. And corresponding quantification of SCI. Scale bar=500μm

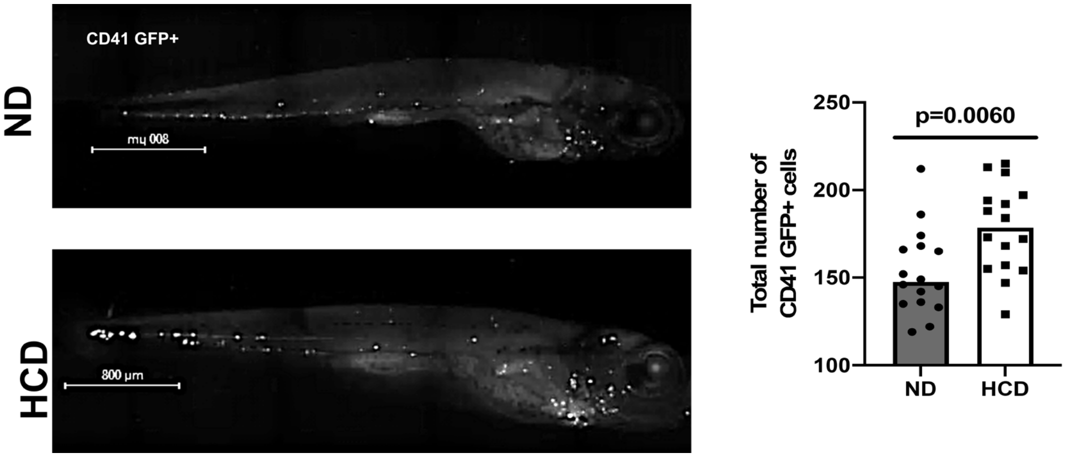


**<u>Suppl. Figure 5:</u>** NASH larvae show increased number of hematopoietic stem cells (**A**) Representative maximum intensity projections of 13 days-post fertilization (dpf) transgenic larvae fed with normal diet (ND) and high cholesterol diet (HCD) at different days of feeding. **(B-C)** Quantification of CD41+ cells in whole larvae.

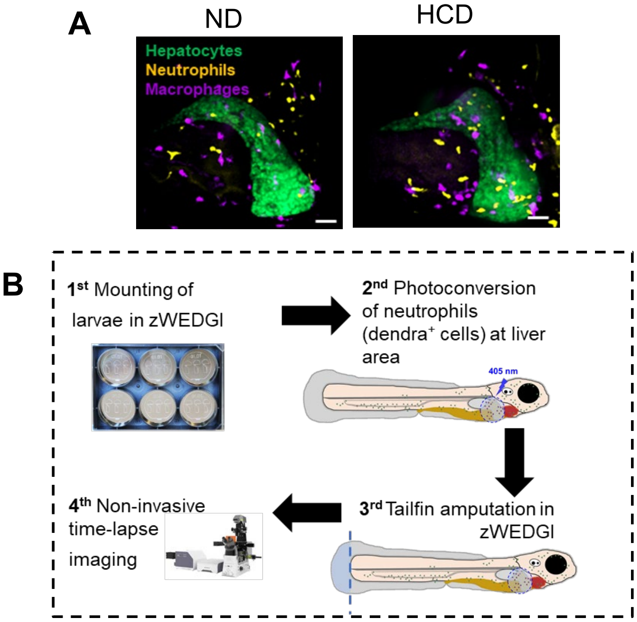


**<u>Suppl. Figure 6:</u>** Neutrophil reverse migration from liver area. (A) representative image of neutrophil recruitment at liver area of larvae fed with Normal diet (ND) and High Cholesterol Diet (HCD). (B) Diagram of experimental set up to evaluate the percentage of neutrophils that reverse migrate from liver area to tailfin transection injury.
